# CrossFilt: A Cross-species Filtering Tool that Eliminates Alignment Bias in Comparative Genomics Studies

**DOI:** 10.1101/2025.06.05.654938

**Authors:** Kenneth A. Barr, Yoav Gilad

## Abstract

Comparative functional genomic studies are often affected by biased read mapping across species due to inter-species differences in genome structure, sequence composition, and annotation quality. We developed CrossFilt, a filtering strategy that retains only sequencing reads that map reciprocally between genomes, ensuring that quantification of read counts is based on directly comparable genomic features. Using both real and simulated RNA-sequencing data from primates, we show that CrossFilt outperforms five alternative approaches that are commonly used, resulting in more accurate inference of gene expression differences. Our results highlight the impact of preprocessing strategies on the analysis of cross-species functional genomics data.

## Background

The field of functional genomics has adopted sequencing-based technologies to measure cellular activity, such as gene expression, chromatin accessibility, methylation, splicing, and other molecular phenotypes. The first step in nearly any sequencing-based functional genomics study is to align sequencing reads to a reference genome to identify their location. In most studies, genomic annotations are used to categorize and contextualize the sequencing read counts. For example, reads that map to annotated coding regions may be counted and compared across samples to identify inter-individual differences in the gene expression levels.

For studies involving samples from the same species, the alignment and annotation steps are relatively straightforward. Although polymorphisms can affect the mapping probability of sequencing reads to specific genomic regions (1), effective approaches exist to address this issue (2), and overall, the sequences of different genomes are highly similar between individuals of the same species. Moreover, functional annotations and context of the mapped reads are, by definition, identical between individuals of the same species, such that even errors in annotation do not bias the results of the comparisons, because they are shared across the entire sample.

Cross-species functional genomics comparisons introduce more nuanced challenges. In addition to the alignment and mapping steps, which must be done for multiple species, one must identify orthologous and syntenic genomic features (3). For example, to meaningfully compare gene expression levels between humans and other primates, one must identify and annotate orthologous genes in the studied species. Once these orthologous and syntenic regions are identified – a task that is by no means trivial – one must also account for different read alignment properties across species. The problem is that if the probability of sequence alignment to annotated features differs between species (for example, because genes have different length in different species), it will introduce bias when comparing estimated quantities (e.g., gene expression levels) across species.

Alignment bias emerges when there is inter-species variation in the length, copy-number, or structural organization of annotated features. A genomic feature that is longer in one species than another, whether due to differences in exon count, repetitive sequences, or expansion of regulatory elements, provides a larger target for mapping sequencing reads. This can result in an artificially higher measured signal, even if the underlying molecular activity is identical. For instance, using RNA-sequencing data, one might observe apparent higher transcript abundance in a species with longer exons, despite no actual difference in gene expression levels across species. Conceptually, correcting for differences in the length of corresponding features across species is straightforward if precise structural information is available. In practice, however, most comparative studies typically define corresponding regions across species without detailed information on potential differences in length and structural organization.

Inter-species differences in genomic duplications also contribute to alignment bias. For example, a chromatin accessibility assay could indicate lower read counts in an accessible region in one species simply because the corresponding enhancer is within a perfectly duplicated region elsewhere in the genome. Some regions exist in a single copy in one species but have duplicates in another. When this occurs, sequencing reads from duplicated regions may be removed from the alignment data due to mapping ambiguity, making it difficult to know whether observed inter-species differences reflect true changes in molecular activity or artifacts of mapping redundancy.

This issue is particularly relevant for gene families that have expanded in certain lineages, as well as for conserved regulatory sequences that have been duplicated and repositioned across the genome (4,5).

Differences in annotation quality between genomes add further complication. Some species, such as human and mouse, have highly curated genome annotations that comprehensively define genes, transcripts, and regulatory elements (6). Other genomes are often annotated using incomplete or lower-confidence annotations. When species-specific annotations are projected from one genome coordinate system to another, researchers may inadvertently compare a well-annotated genome to one with missing features, mistaking absent annotations for true biological differences. This issue is particularly problematic for regulatory sequences, which are often annotated based on indirect evidence (such as sequence conservation or chromatin state) and may be underrepresented in species with less extensive functional genomics data.

To address these challenges, we developed CrossFilt (Cross-species Filtering Tool), an alignment-based method that mitigates mapping biases by excluding sequencing reads that do not map reciprocally between species. In this study, we applied CrossFilt to both real and simulated RNA-sequencing data from humans, chimpanzees, and rhesus macaques (7) to identify differentially expressed genes between the species. Our aim was to assess how different alignment and feature definition strategies influence the outcomes of comparative genomics analyses.

Compared to five alternative methods commonly used for cross-species sequencing data, CrossFilt reduced the number of false positives and improved the consistency of inferred gene expression differences between species. By restricting analysis to reciprocally mappable reads, CrossFilt ensures that feature quantification is based only on directly comparable genomic regions. More broadly, we demonstrate that the methodological choices made during data processing can substantially affect the set of molecular differences identified between species, with some existing and commonly used approaches producing markedly inflated false positive rates.

## Results

### CrossFilt, a novel read filtering strategy for comparative genomic studies

CrossFilt is a cross-species filtering approach that is designed to facilitate unbiased comparisons of sequencing-based data obtained from different species. It uses a reciprocal liftover (8) strategy to retain only the sequencing reads that map accurately and consistently to the genomes of each species. In the following vignette, we describe CrossFilt as applied to a typical case: a pairwise comparison of RNA-sequencing data between two species (e.g., human and chimpanzee). Here, the ‘target’ and ‘query’ genomes respectively correspond to the human and chimpanzee reference genomes. CrossFilt also requires genome annotations for each species but is agnostic to their source, allowing flexible integration with any annotation set of choice.

CrossFilt’s reciprocal liftover approach relies on chained alignments between species, which are needed to accurately convert coordinates from the genome of one species the other. A chain is a sequence of pairwise alignments that are arranged in a single order and orientation; by definition, this arrangement must be consistent with the order of genomic sequence in both species (thus, chains include neither translocations, duplications, nor inversions). The CrossFilt process begins by aligning sequencing reads derived from the target samples to the target reference genome. For each aligned read, CrossFilt uses the reference genome coordinate system to identify its start and end positions. It then retrieves all chains that fully encompass the read (see Methods). If more than one chain fully encompasses a single read, CrossFilt selects the chain with the highest UCSC alignment score (9), which reflects the length and quality of the alignment, favoring long, gap-free regions with few mismatches. Next, each nucleotide in the read that matches the target genome is replaced with the corresponding nucleotide from the query genome, accounting for insertions and deletions. CrossFilt excludes reads if their start or end coordinates coincide with insertions or deletions in the query genome.

The lifted reads are then converted into FASTQ format and independently realigned to the query genome using the query species’ annotations. A custom script (*crossfilt-filter*; see Supplementary Materials) verifies that the realigned reads match exactly with the lifted BAM alignment in terms of start position, read name, and concise idiosyncratic gapped alignment report (CIGAR) string. Reads that fail this verification are discarded. The reads passing this step are reciprocally lifted back to the target genome, and the script *crossfilt-filter* is used again to confirm that each read maps precisely to its original genomic location. Only reads successfully passing both reciprocal liftover steps are retained.

The next step is to define orthologous features between the two species. As mentioned, CrossFilt is compatible with any number of methods for orthology inference, whether based on sequence identity, predictive models, or external data. As our study includes data from humans, chimpanzees, and rhesus macaques, we elected to use the Comparative Annotation Toolkit (CAT, (8)), which provides comprehensive annotations for all three species (see Methods). After separately mapping reads from each species to the orthologous features in the corresponding reference genome, CrossFilt keeps only the reads that map consistently to the same genomic feature in both species. Reads mapping inconsistently between species are removed. Through this rigorous reciprocal filtering process, CrossFilt effectively mitigates alignment and annotation biases, ensuring robust comparative analyses.

### Commonly used strategies for comparing sequencing-based data across species

Comparative functional genomics studies have used different strategies to define orthologous features and quantify molecular phenotypes across species using sequencing-based data. These strategies differ in three key respects: (***i***) whether sequencing reads from all species are aligned to a single reference genome or to species-specific genomes, (***ii***) how functional annotations are assigned to define genomic features such as enhancers, genes, or exons in each genome, and (***iii***) whether steps are taken to correct for read mapping biases introduced by sequence divergence, structural variation, or differences in annotation quality across genomes of different species.

Approaches that align all reads to a single genome bypass the need to define orthologous features explicitly but are prone to alignment bias when mapping efficiency is affected by sequence divergence between the species (e.g., when chimpanzee-derived reads are aligned to the human genome). In contrast, approaches that align reads to the corresponding genome of each species (e.g., reads from human samples are aligned to the human genome, and reads from the chimpanzee samples are aligned to the chimpanzee genome) require definition of orthologous sequences or functional features, either by name matching or via formal annotation transfer methods. Additional bias correction strategies such as read- or feature-level filtering, further distinguish methods in terms of their robustness and reliability (as we show in our analysis below).

We compared CrossFilt to five commonly used approaches that span these design choices. We refer to these approaches as single-reference, consensus, dual-reference, annotation transfer, and orthologous exons. Below we briefly describe each approach and highlight their primary differences. A comparison of relevant features of these approaches is available in **Table 1**.

**Table 1:**
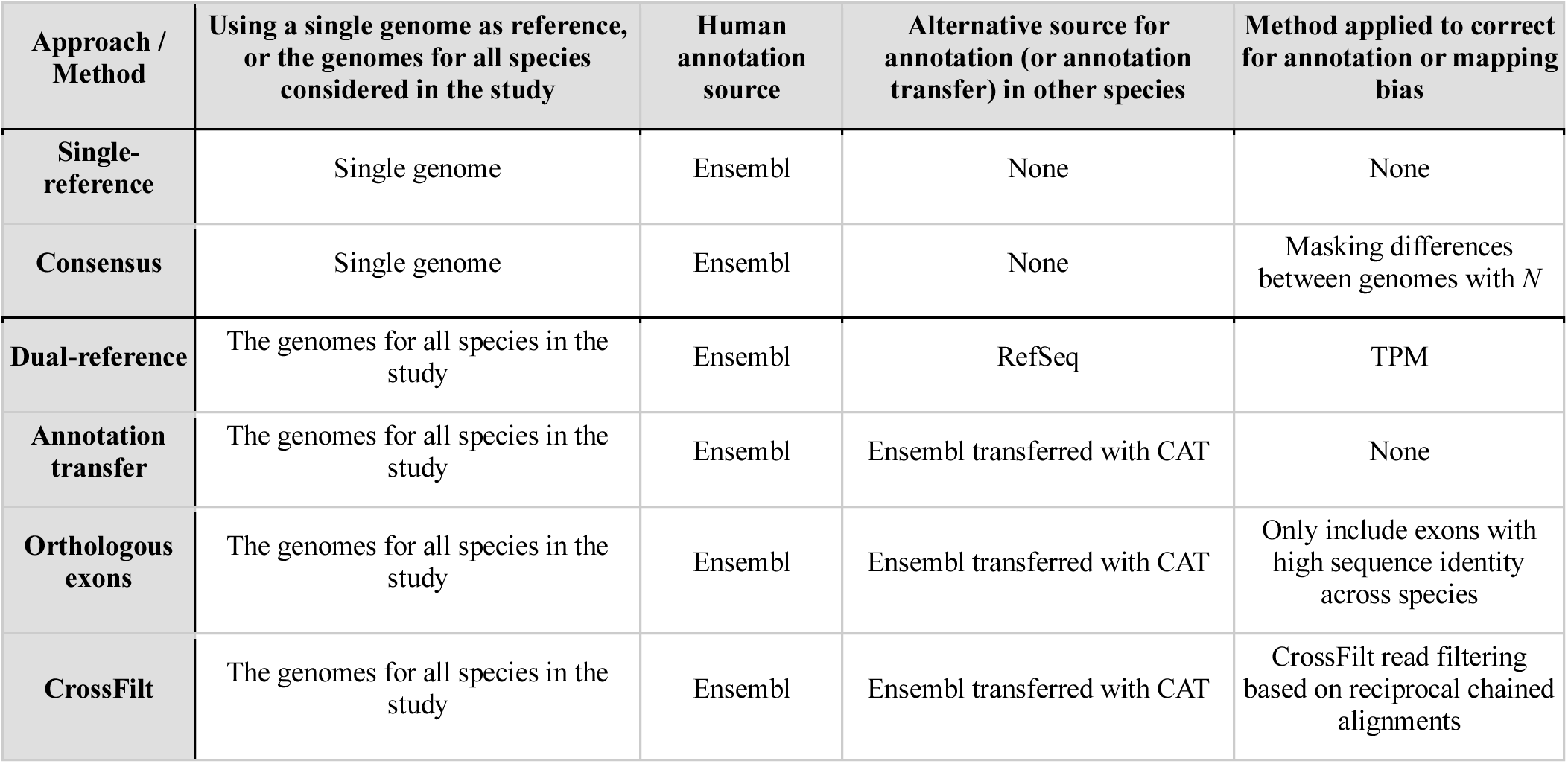
Summary of compared approaches.

#### Single-reference

In this approach, used by (11) to build a single-cell atlas of primate brain development, sequencing reads from multiple species are aligned to the genome of a single species, and the corresponding functional annotation (e.g., Ensembl (12)) is used. No attempt is made to correct for mapping bias or annotation mismatches. This approach is straightforward and does not require ortholog definitions, but the alignment is sensitive to sequence divergence between species.

#### Consensus

Used by (11,13,14), a synthetic reference genome is created by masking all nucleotide differences between the studied species with ‘N’s, to create a reference genomic sequence. Reads from all species are then aligned to this masked consensus genome using a single genomic annotation set. This removes alignment bias at divergent sites but does not address broader structural differences or annotation mismatches. The use of a single annotation also avoids the need to explicitly define orthologous features.

#### Dual-reference

In this approach, used by (15–24) and others, reads from each species are aligned to their respective reference genomes. In a comparative study involving human and chimpanzee samples, for example, it would be reasonable to use the best available annotation set for each species, even if they differ in terms of gene structure and completeness. For instance, human reads could be aligned to the Ensembl-annotated human genome, while chimpanzee reads could be aligned to the panTro6 assembly using RefSeq annotations. Transcripts Per Million (TPM) normalization is typically applied to normalize gene expression levels, as this accounts for differences in gene length between samples. Orthology can then be defined implicitly, by matching gene names across species. This approach is simple but may be biased when annotation quality differs substantially between species.

#### Annotation transfer

Used by (25–28) and others, annotations from one genome are transferred to the other species using genome alignment tools such as CAT (10) or TOGA (Tool to Infer Orthologs from Genomic Alignments) (29). This produces consistent gene models across species and avoids the limitations of low-quality species-specific annotations. However, no further steps are taken to address alignment bias, so mapping artifacts may still contribute to false positives.

#### Orthologous exons

Several studies, including from our group, have used a more conservative variant of annotation transfer (30–35) in order to reduce alignment and annotation bias. In the ‘orthologous exons’ approach, exons are filtered after the annotation transfer step to ensure that only highly comparable features are included in the analysis. For example, one might retain only the exons with high sequence identity (e.g., > 85% for a comparison of human and chimpanzee) and minimal length differences (e.g., < 10%) between species.

#### CrossFilt

Our method, CrossFilt, uses a reciprocal alignment strategy to filter reads prior to quantification. Reads are retained only if they map precisely and consistently to both species’ genomes after accounting for insertions, deletions, and mismatches. This is combined with ortholog definitions derived from annotation transfer (e.g., CAT annotations). By filtering at the read level and not just at the feature level, CrossFilt minimizes both alignment and annotation bias, yielding reliable cross-species comparisons.

### How did we compare CrossFilt with other approaches?

To systematically compare these approaches against CrossFilt, we evaluated all six methods in the context of a comparative gene expression study using RNA sequencing data from human, chimpanzee, and rhesus macaque. Our goal was to assess the performance of each approach not only in terms of the number of genes tested, but also the quality and reliability of the resulting inferences, focusing on identifying differentially expressed genes between the species. While maximizing gene inclusion is desirable, doing so without accounting for differences in annotation quality or mapping efficiency may introduce bias. The most effective methods are therefore those that strike an optimal balance between inclusiveness and accuracy.

We structured our comparison around three distinct analyses. We began by assessing the extent and symmetry of genome and transcriptome coverage across species for each method. These metrics reflect both the number of genes that can be reliably tested and the completeness of their annotations. For example, an annotation set with more transcripts in one species than another may lead to asymmetric read mapping, potentially biasing estimates of gene expression (because when using RNA sequencing data, gene expression is a relative measurement (36,37)). We evaluated coverage in two ways: First, by comparing the number of genes that passed method-specific filtering criteria and would be included in each pipeline. Each method specifies different criteria for inclusion. For example, the ‘single-reference’ method is the most inclusive in this context, as it uses all annotated human genes for the comparison. Other methods include filtering criteria such as ‘minimum sequence identity’ or ‘length similarity’ (required by the ‘orthologous exons’ method), or ‘reciprocal mapping’ (required by CrossFilt). Second, we measured the degree of annotation completeness across species, using the proportion of each gene’s sequence that can be covered by aligned RNA-sequencing reads. For example, in the ‘consensus’ approach, the number of ‘N’ sequence replacements in certain genomic regions is large enough to exclude unique read mapping to these regions.

Next, to evaluate the accuracy and robustness of each method, we simulated RNA-sequencing data with known inter-species differences in gene expression. This allowed us to directly measure statistical power and false discovery rates across approaches, as well as to detect alignment and mapping induced species-specific biases in apparent gene expression levels or effect size estimates. We performed differential expression analysis and compared the proportion of true and false positives identified using each method. We assessed how well each method recovered the simulated inter-species differences in gene expression, and quantified bias in the direction and magnitude of false positive signals. This analysis also allowed us to evaluate whether errors were systematically enriched in genes with low expression or genes duplicated in one species.

Finally, we applied each method to real data, using previously collected bulk RNA-sequencing data from human, chimpanzee, and macaque tissues (7). Using consistent quality control and normalization procedures across all pipelines, we identified differentially expressed genes in pairwise species comparisons and evaluated consistency across methods (because, using real data, we do not have a gold standard set of differentially expressed genes). Specifically, we assessed the overlap in differentially expressed gene sets, the concordance of effect size estimates, and the inferred sharing of inter-species differentially expressed genes across tissues. We also performed gene set enrichment analysis to determine whether methodological differences influenced downstream biological interpretation. By examining patterns of gene sharing, effect size correlation, and enrichment among functional categories, we evaluated how methodological choices impact the interpretation of interspecies differences.

This three-part evaluation framework allowed us to compare the strengths and limitations of each approach and identify sources of bias that may distort downstream analyses. In the sections that follow, we present the results of these comparisons and highlight the distinct advantages of CrossFilt relative to existing approaches.

### Comparison of genomic and genic coverage across approaches

The number of genes retained for differential expression analysis varied substantially across approaches (**Figure 1A-B; Table S1**). The ‘single-reference’ and ‘consensus’ approaches use complete, unfiltered human genome annotations. The ‘single-reference’ approach thus includes nearly all genes annotated in the human genome (38,584 genes). The ‘consensus’ approach includes slightly fewer genes (**Figure 1C**), with the exact number depending on the threshold used to define a gene’s ‘mappability’. For example, if we use 50% sequence masking as the cutoff beyond which of a gene is considered unmappable, 316 and 1,163 genes would be excluded from the human-chimpanzee and human-macaque comparisons, respectively. These masked regions reflect sites of sequence mismatch (divergence), insertion, or deletion between species, and are therefore expected to contribute to alignment bias in the ‘single-reference’ approaches. As expected, given the evolutionary distance between species, we found that genes in the human-macaque consensus genome contains many more masked positions (14.7%) than the human–chimpanzee genome (3.6%). This observation explains why divergence-driven alignment bias is expected to be more severe in more distantly related species.

**Figure 1:**
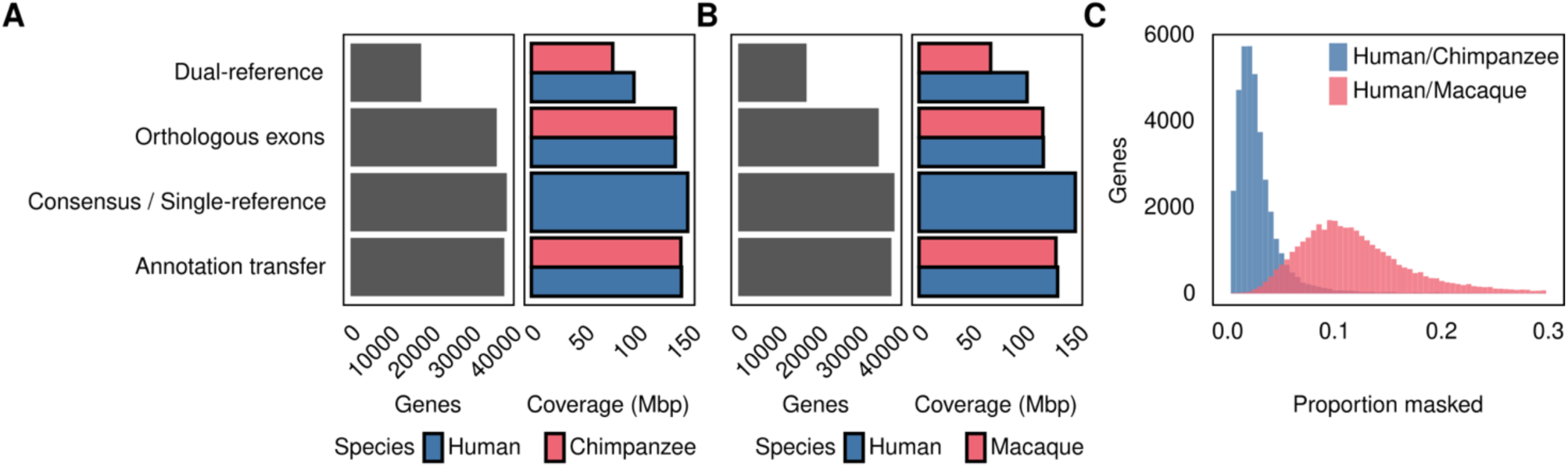
Genomic and gene coverage. **(A)** The number of genes and megabases of genomic coverage for each annotation approach when comparing human and chimpanzee RNA expression. **(B)** The number of genes and megabases of genomic coverage when comparing human and macaque RNA expression. **(C)** Histogram of the proportion of each gene masked by ‘N’ when using a consensus genome for the indicated species comparisons, excluding genes with greater than 30% masking (488 genes from the human-chimpanzee comparison and 2,575 from the human-macaque comparison).

The ‘dual reference’ approach resulted in the smallest set of comparable genes (17,443 in the human-chimpanzee comparison and 16,885 in the human-macaque comparison) due to the more limited annotations available for the chimpanzee and macaque genomes. We also observed substantial inter-species differences in read coverage under this approach (**Figure 1A-B**). Notably, the ‘dual reference’ method differs from the others in that orthologous genes are inferred by matching gene names between species. As a result, it retains primarily protein-coding genes, since other transcripts often lack standardized gene names across species and are excluded. The other approaches (‘annotation transfer’, ‘orthologous exons’, and CrossFilt) use sequence-based definitions of orthology that do not depend on gene names. The ‘annotation transfer’ and ‘orthologous exons’ methods performed better in terms of gene inclusion, though the ‘orthologous exon’ approach excluded a larger number of genes due to its strict sequence similarity filters (**Table S1**). In our implementation of CrossFilt, we used CAT for comparative annotations, which resulted in identical gene counts as ‘annotation transfer’.

### Assessing differential expression accuracy using simulated data

To evaluate how alignment and annotation choices influence differential expression (DE) analysis, we simulated bulk RNA-sequencing data from human, chimpanzee, and rhesus macaque tissues (heart, liver, lung, and kidney). Using gene-specific expression means and dispersion estimates derived from empirical data we collected in a previous study (7), we simulated the expression of 34,693 genes in humans and chimpanzees and 31,807 genes in humans and macaques in four biological replicates per species and tissue combination (see Methods for details). In each pairwise species comparison, we randomly selected 10% of genes to be differentially expressed, with half showing higher expression in human and the other half in the comparison species (simulated inter-species log fold-changes were set to 2). We then applied each of the six approaches to align and process the simulated reads, followed by DE testing using a standardized limma-voom framework (see Methods). When we used the ‘dual-reference’ method, we applied TPM normalization instead of counts-per-million (CPM) to account for gene length differences between species.

With respect to false positive error rate, all six methods performed better in the human–chimpanzee comparison than in the more divergent human–macaque comparison (**Figure 2A-B, Table S2**) CrossFilt exhibited the strongest overall performance, achieving the lowest empirical false discovery rate (eFDR of 4.0% in both species comparisons) without a reduction in power (**Figure 2C**). The ‘consensus’ approach ranked second in accuracy by these metrics (eFDR of 5.4% and 8.9% in the comparisons of human-chimpanzee and human-macaque, respectively). Among the remaining approaches, performance varied depending on the comparison. For example, in the human–chimpanzee comparison, the ‘single-reference’ approach produced fewer false positives (eFDR of 6.8%) than the ‘annotation transfer’ or ‘orthologous exons’ methods (eFDR of 12.0% and 10.9%, respectively). However, in the human-macaque comparison, the ‘single-reference’ approach resulted a marked increase in false positives (eFDR of 18.5%). The ‘dual reference’ approach, which is commonly used in comparative genomics studies in primates (15–24), resulted the worst performance in both comparisons, with eFDR values of 53% for human-chimpanzee and 57% for human-macaque.

**Figure 2:**
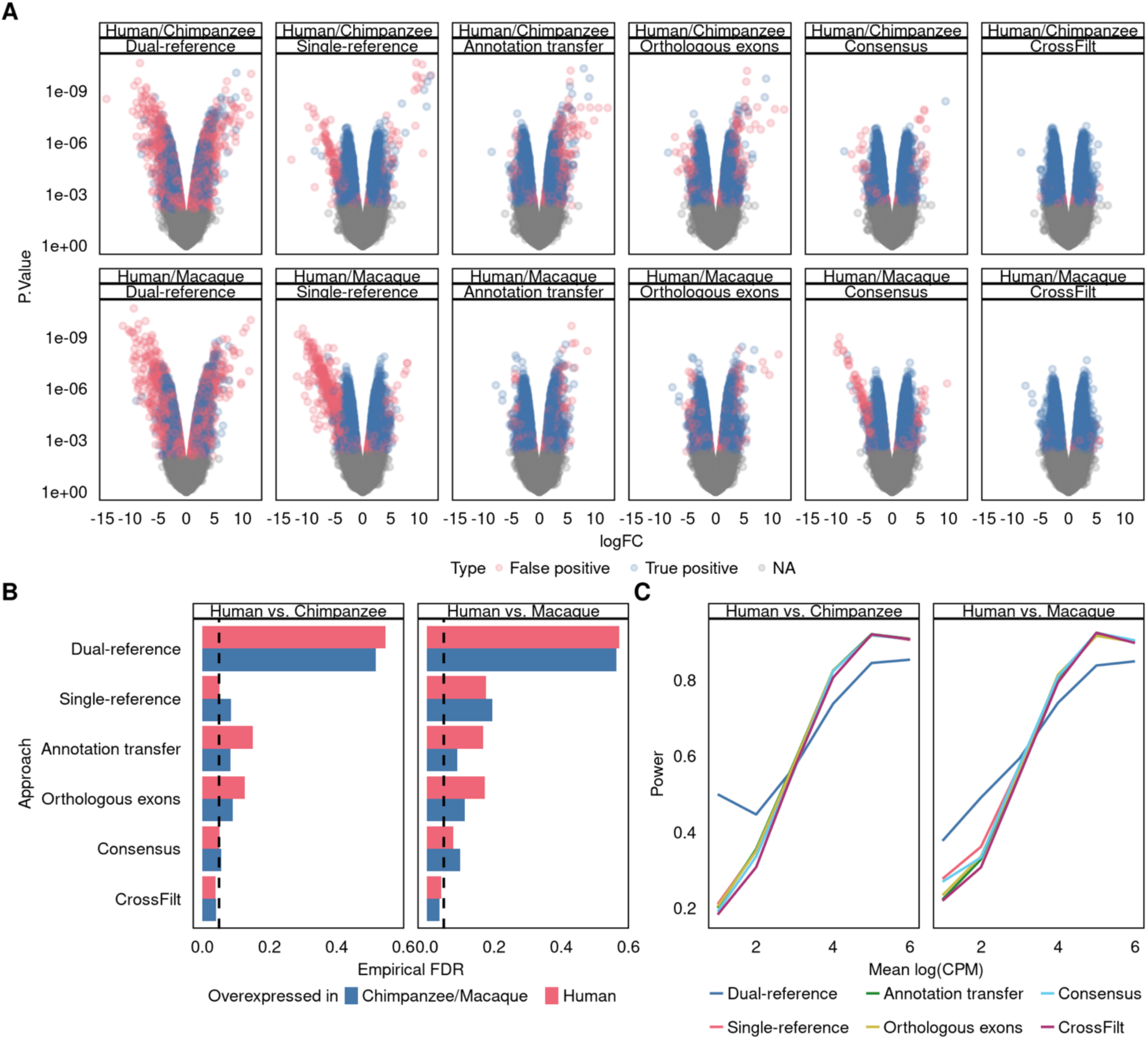
Simulation performance. **(A)** Volcano plots for each comparison and approach. Genes with FDR ≤ 5% are highlighted in red (false positives) and blue (true positives). Positive values of logFC indicate genes overexpressed in humans while negative values indicate genes overexpressed in the comparison species. **(B)** The empirical FDR for each approach, split depending on the species in which the gene has higher expression. **(C)** Power to detect true positives in each approach as a function of mean expression across species.

We evaluated the specific properties of false positives across approaches. Genes falsely identified as DE tend to have lower average simulated expression levels in the ‘single-reference’ and ‘consensus’ approaches compared to all other approaches (*P* < 1.3 x 10^-5^ in all pairwise comparisons; **Figure S1**). Additionally, we observed species-specific asymmetry in the number and effect sizes of false positives. When using the ‘single-reference’ and ‘consensus’ pipelines, we identified fewer false positives classified as more highly expressed in human (**Figure 2B**). These false positives tend to have smaller effect sizes and higher expression levels relative to false positives classified as more highly expressed in chimpanzees or macaques (**Figure S2**). In contrast, for the other approaches, we identified more false positives with higher expression in humans. These false positives tend to have larger effect sizes and overall lower expression levels. The degree of species asymmetry in false positives was greatest for the ‘annotation-transfer’ method (67% of false positives associated with higher expression in human in the pairwise comparison with each species), followed by ‘orthologous-exons’, ‘single-reference’, ‘consensus’, and ‘dual-reference’ (**Table S2**). The DE results using CrossFilt do not show such bias.

Taken together, our simulation results highlight the susceptibility of some methods, particularly those that do not correct for alignment bias, to elevated false discovery rates and systematic distortions in estimated effect sizes. CrossFilt consistently showed the best overall performance, suggesting that reciprocal read filtering substantially improves the reliability of DE inference in cross-species sequencing-based comparisons.

### Assessing differential expression accuracy using real RNA sequencing data

To evaluate how methodological differences affect downstream biological conclusions, we also applied all six pipelines to real bulk RNA-sequencing data from human, chimpanzee, and rhesus macaque tissues (7). Following our approach for analyzing the simulated data, each pipeline incorporated consistent quality control and normalization steps (see Methods and Supplementary Information); however, in this case, we do not have a gold standard comparison. Therefore, to assess the concordance of DE results, we first examined the effect sizes of genes identified as DE using at least one approach (at FDR ≤ 5%). We compared these DE genes to those identified using all other approaches, considering each inter-species and inter-tissue comparison independently. In general, effect size estimates were consistent across approaches, with DE genes clustering primarily by tissue (**Figure 3A**, **Figure S3**). Consistent with the results from the simulated data, the ‘dual-reference’ approach resulted in the least agreement with the other pipelines (mean R = 0.48 vs. mean R = 0.79 for all other approaches). We ruled out TPM normalization as the cause of this discrepancy (**Figure S4**) and concluded that alignment and annotation asymmetries specific to the ‘dual-reference’ method contribute to this pattern. This effect was most pronounced in the human–chimpanzee comparison, where all other approaches perform more similarly.

**Figure 3:**
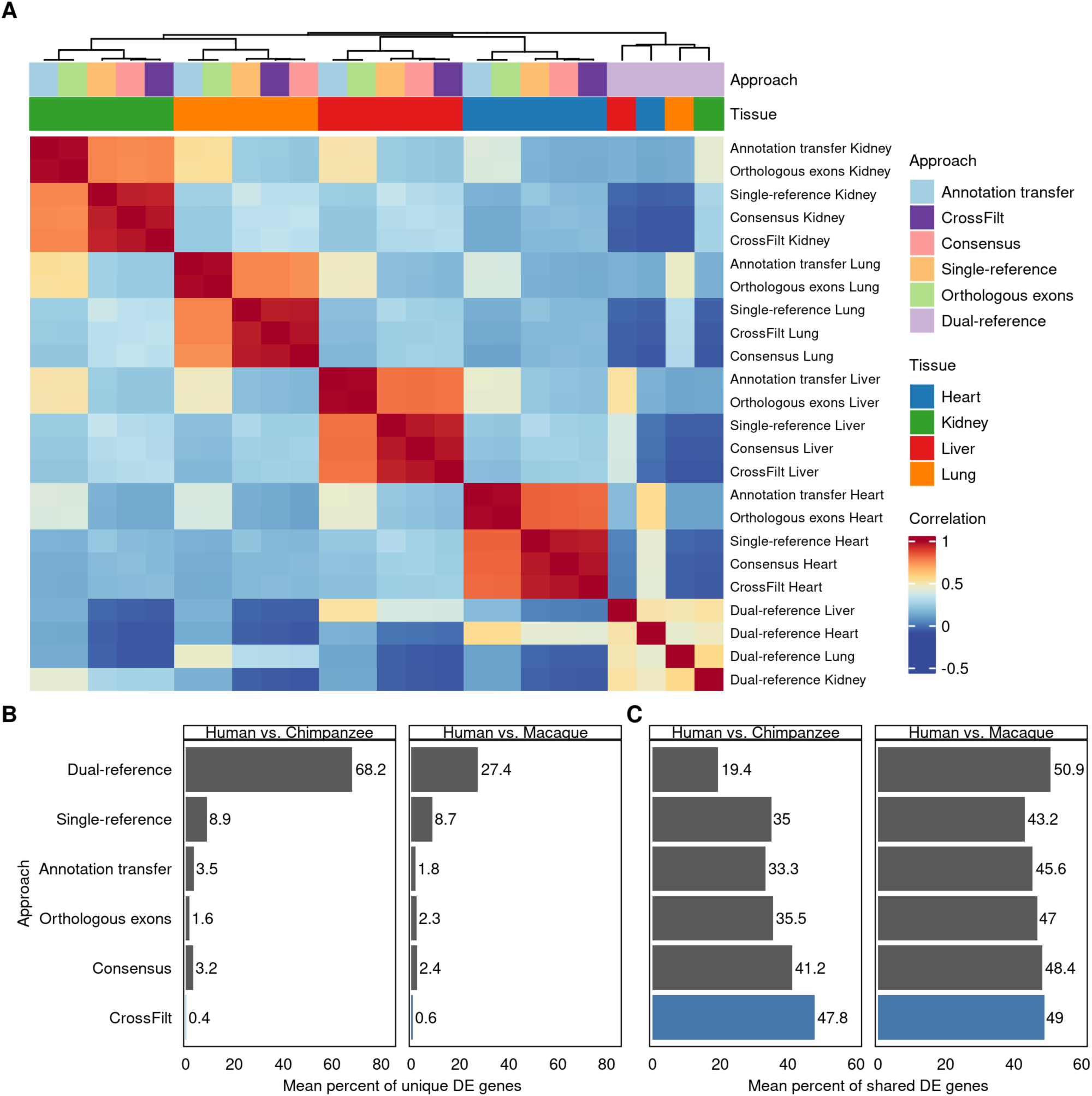
Concordance of DE gene sets in real data. **(A)** Heatmap of the Pearson correlation in human-chimpanzee effect size estimates for genes detected as DE in at least approach and tissue. Order of rows and columns is set by hierarchical clustering. **(B)** The percent of DE genes identified through each approach that are classified as DE by any other approach. **(C)** The percent of DE genes identified through each approach that are also classified as DE by all other approaches.

Across tissues, 68.2% of inter-species DE genes identified by the ‘dual-reference’ pipeline were not identified as DE by any other method (**Figure 3B**). Given the results of the analysis of the simulated data, we considered the DE genes identified only when the ‘dual-reference’ pipeline is used to be false positives. Although the proportion of false positives was lower in the human–macaque comparisons relative to the human-chimpanzee comparisons, the ‘dual-reference’ false positive rate was always higher than for any of the other methods. When we considered a comparison of DE genes identified using all other approaches, we noted that, overall, fewer than half of the DE genes were identified by every approach (**Figure 3C**), highlighting the profound impact of the choice of alignment and annotation strategy on DE inference.

We next used mash (38) to estimate the extent of DE gene sharing across tissues. Because alignment bias should be independent of tissue type, approaches affected by such bias are expected to overestimate the number of interspecies DE genes shared across tissues. Consistent with this expectation, the ‘dual-reference’ approach resulted in the identification of the highest proportion (75.3% in human-chimpanzee and 27.9% in human-macaque) of inter-species DE genes shared across tissues (**Figure 4A**). Using CrossFilt resulted in the lowest estimate (52.4% in human-chimpanzee and 18.7% in human-macaque) of shared interspecies DE genes across tissues. In general, genes identified as interspecies DE in multiple tissues, but not by CrossFilt, were highly enriched for genes identified as false-positives in our simulated data (**Figure 4B**).

**Figure 4:**
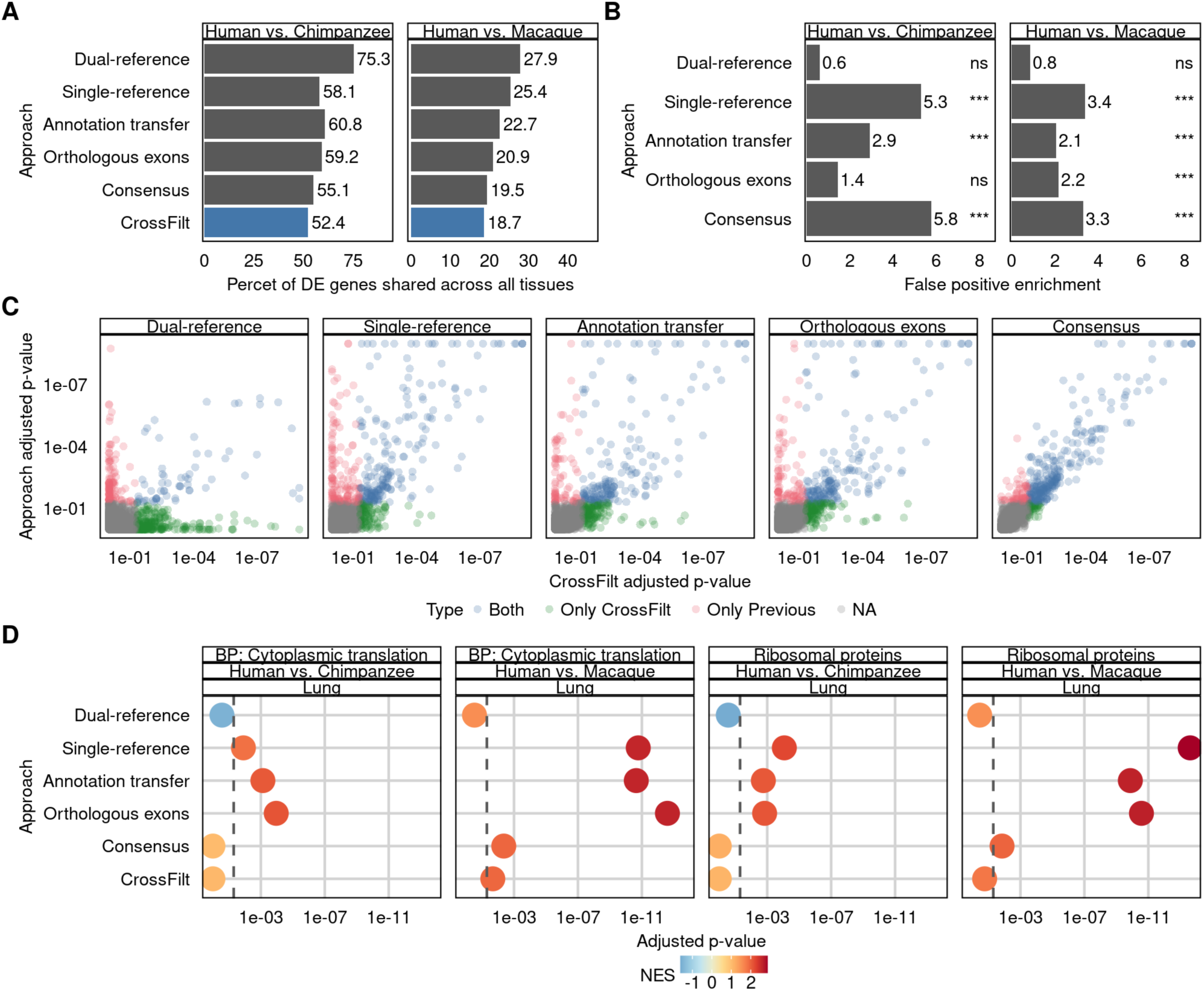
Cross-tissue sharing of DE and gene-set enrichment in real data. **(A)** The percent of DE genes with shared effects across all four tissues, estimated by MASH. **(B)** Enrichment of false positives in simulated data for genes that are identified as shared by the indicated method, but not CrossFilt, relative to genes identified as shared by both approaches. P-values are from one-sided Fisher’s exact test (ns = not significant; *** = P < 0.001). **(C)** Gene-set enrichment (GSEA) significance of Gene Ontology: Biological Process terms from the indicated approach (y-axis) and CrossFilt (x-axis). Terms are colored based on whether the term was significant in only the indicated approach (red), only CrossFilt (green) or both (blue). **(D)** GSEA significance of the term ‘Cytoplasmic translation’ and ribosomal proteins (RPS and RPL genes) in lung. Points are colored based on normalized enrichment score (NES).

Finally, we assessed the extent to which methodological differences affect downstream biological interpretations by performing gene set enrichment analysis. Using ranked t-statistics for each method, we tested for enrichment of DE genes among Gene Ontology biological process categories. While most gene sets produced broadly concordant enrichment scores across pipelines, we identified cases where alignment artifacts may have inflated statistical significance (**Figure 4C**). For example, the category ‘cytoplasmic translation’ was significantly enriched among genes identified as inter-species DE using the ‘single-reference’, ‘annotation-transfer’, and ‘orthologous-exons’ pipelines. Further inspection revealed that this enrichment was driven by a group of ribosomal genes with high sequence similarity, likely resulting from ambiguous read mapping and artifacts in gene-level quantification (**Figure 4D**).

Taken together, these analyses underscore that methodological choices influence not only which genes are identified as differentially expressed, but also how consistent those results are across contexts (in our case, tissues) and how they are interpreted in a functional context. CrossFilt’s design minimizes alignment and annotation bias and avoids many of the artifacts observed in alternative methods.

## Discussion

Comparative functional genomics studies have consistently faced the challenge of alignment bias. Whether subtle or pronounced, alignment bias can subsequently affect nearly every step of sequence-based quantification across species. Despite widespread recognition of this problem and many attempts to mitigate its impact (including our prior work using the ‘orthologous exons’ approach (39)), residual bias remains a persistent source of false positives identification of interspecies differences, and distorted biological interpretations. As our results demonstrate, CrossFilt substantially improves the accuracy of cross-species quantification by removing sequencing reads that cannot be mapped unambiguously between species.

While some comparative studies have adopted methods to reduce alignment artifacts, many continue to rely on pipelines that are fundamentally flawed. ‘Single-reference’ and ‘dual-reference’ approaches, for example, remain in use despite their susceptibility to a high degree of alignment and annotation asymmetries. These approaches tend to inflate expression differences where none exist, particularly when genome annotations differ in quality or completeness between species. There are also many comparative studies that addressed this problem through targeted feature filtering (limiting analyses to orthologous exons with high sequence similarity is one such example), but as we show, CrossFilt offers a more comprehensive and general solution by addressing the issue directly at the level of read mapping.

The importance of resolving alignment bias extends beyond generating a more accurate list of differentially expressed genes between species. The primary value of comparative functional genomics lies in its ability to identify general biological patterns and molecular mechanisms that distinguish species, cell types, or environmental responses. Biases in read alignment and feature annotation can systematically distort this inference, leading to incorrect conclusions about how molecular phenotypes differ across species. For example, in studies of tissue-specific expression, apparent inter-species similarities in gene regulation across tissues may be overestimated because alignment artifacts will be common to the analysis of data from all tissues. Similarly, when comparing gene expression responses to treatment, diet, or environmental exposures, biased mapping can obscure or exaggerate shared species-specific effects and prevent us from finding exposure- or treatment-specific patterns. In all of these contexts, correcting for alignment bias is necessary not only for precision, but for biological validity. Furthermore, functional insights based on gene ontology or pathway enrichment analysis may also be susceptible to alignment bias, because gene function is often correlated with evolutionary constraint. As a result, divergence, and therefore the degree of mapping bias, can vary systematically across functional categories, potentially leading to spurious enrichment signals or the masking of true biological patterns.

The problem is not specific to gene expression. Although we chose RNA sequencing data as the motivating example, and genes as the primary features of interest, the same limitations apply to any sequencing-based molecular phenotype used in a comparative framework. Whether the goal is to compare transcription factor binding, chromatin accessibility, splicing, or other read-based measurements across species, differences in sequence composition, structure, or annotation can lead to asymmetric mapping and biased quantification. CrossFilt addresses this issue in a general way. It does not depend on a particular annotation strategy and can be applied to any analysis involving orthologous genomic features as long as genome sequences and alignment chains are available.

Because CrossFilt operates at the read level and is annotation-agnostic, it can be used flexibly with any method of ortholog definition, including curated annotations, projection tools, or inference-based pipelines. It is compatible with both protein-coding and noncoding features, and readily integrates into existing differential analysis workflows. While our evaluation focused on inter-species comparisons among primates, the method can be extended to other taxa, including more divergent species, where annotation asymmetries and mapping biases are often even more pronounced.

Finally, CrossFilt provides a conceptual framework for improving data quality in comparative genomics. As functional genomics expands to incorporate more species, cell types, and molecular phenotypes, the demand for robust, bias-aware preprocessing methods will continue to grow. CrossFilt can help ensure that downstream comparisons are meaningful, that differences reflect biology rather than technical artifacts, and that conclusions drawn from cross-species data are genuinely comparative in nature.

## Conclusion

Our findings underscore the importance of methodological choices in comparative genomics. Without careful control for genome assembly and annotation differences, comparisons based on sequencing data can be misleading, attributing biological significance to technical artifacts. By providing a robust and scalable approach for cross-species analysis, CrossFilt enhances the accuracy of comparative genomics and helps clarify the true molecular differences between species.

## Methods

### RNA-sequencing data

We previously collected 50 bp single-end RNA-sequencing data from human (*Homo sapiens*, all of reported Caucasian ancestry), chimpanzee (*Pan troglodytes*), and Indian rhesus macaque (*Macaca mulatta*) tissues (heart left ventricle, kidney cortex, liver, and lung), which we used to comparatively analyze gene regulation between the species (7). The data include 48 samples in total, with four individuals per species and four tissues per individual; samples were pooled and sequenced on 26 lanes of four different flow-cells on the Illumina HiSeq 2500 platform (7). To prepare the data for the current study, we downloaded FASTQ files from the Sequence Read Archive (accession numbers SRR6900765 through SRR6900812) and used TrimGalore (v0.6.10; https://www.bioinformatics.babraham.ac.uk/projects/trim_galore/), a wrapper based on Cutadapt (v2.6; (40)) to remove sequencing adapters from the reads. We trimmed using a specificity of 3.

### Simulated RNA-sequencing data

We simulated RNA-sequencing data to match the structure of the empirical data (7). We used the mean expression level for each gene in each tissue in the human dataset as a baseline for simulated gene expression levels in both species. We then fit dispersion estimates for every gene using the `estimateDisp` function in edgeR (41,42). We simulated differential expression for 10% of genes. For 5% of the genes, we increased the mean expression level in humans by factor of 4 (corresponding to a log fold change of 2). For the other 5%, we increased expression in the alternate species by a factor of 4. We then used mean and dispersion estimates to generate data for each species and tissue type. Using the `rnbinom` function in R, we generated four biological replicates for each species under a negative binomial model. We then simulated reads from the longest orthologous transcript each of these genes (defined by CAR) using the polyester packaged in R (43).

### Reference genomes and annotation sets

We used the hg38, panTro6, and rheMac10 genome builds from the UCSC genome browser (44) as reference genomes for each species. For each pipeline, we performed genome alignments using STAR (45). We built each reference with the `sjdbOverhang` parameter set to 49 to match the 50bp read-length in our data. For the human annotation set, we used the 10X Genomics Cellranger 2024 GEX annotation set (built from Ensembl v110) (12). Annotation sets for chimpanzee and rhesus macaque vary according to the approach used, as described below (see also **Table** S3). We aligned reads to each of these STAR references using the flags `--outSAMmultNmax 1` and `--outFilterMultimapNmax 1`. We also used `--quantMode GeneCounts` to generate read-counts for each gene.

### CrossFilt

The core functionality of CrossFilt is a custom script (`crossfilt-lift`; available at https://github.com/kennethabarr/CrossFilt) that will lift over a target BAM file and return the sequence and coordinates for each read in the query assembly. Here, the human genome is the target assembly and the alternative genome is the query assembly. For each read, we identify the start and end coordinates of the read and return all chains that fully encompass that read. Then, for every nucleotide in the read that matches the target genome, we replace that nucleotide with the query sequence in that chain, including any insertions or deletions. We reject any reads whose start or end coordinate is an insertion or deletion in the query genome, as well as any reads with insertions larger than 300 bp. If there are insertions, we use the quality of the last sequenced nucleotide to fill missing data in the quality string. We update strand information and CIGAR strings to reflect the new coordinates in the query genome. If a read successfully maps in multiple chains, we only return the read from the highest-scoring chain. In order to filter reads from the target genome alignment, we first lift the reads to the query genome. We then convert the lifted bam to FASTQ format using SAMtools (46) and we realign the lifted reads in the query genome. We then use another custom script (`crossfilt-filter`) to check that these aligned reads have the same reference start, name, and CIGAR string as they did in the lifted BAM alignment. We exclude all reads that either failed to lift or did not align to the same location. We then use `crossfilt-lift` to lift these reads back to the target genome and we verify they returned the original aligned location using `crossfilt-filter`. The previous steps yield BAM files with the reads aligned in the target genome as well as the equivalent reads in the query genome. Using the Comparative Annotation Toolkit (CAT) (10), we find orthologous features and count reads that map to features in both genomes. Reads that do not map to the same feature in both genomes are removed.

### Consensus genome

We used custom scripts to construct human-chimpanzee and human-macaque consensus genomes from the RefSeq genome builds (available at https://github.com/kennethabarr/ConsensusGenomeTools). Starting from hg38, we scanned through the UCSC chain file for the other species and masked each mismatched base with an ‘N’. We also masked six bases on each side of all insertions and deletions. We annotated genes in each species using the human annotations.

### Annotation transfer

We used CAT implementations of TransMap (48) and AUGUSTUS (49) to transfer human annotations to the chimpanzee and rhesus macaque genomes. As input, we used genome alignments obtained from the Zoonomia project’s 447-way mammalian alignment (50). The chimpanzee alignments did not include mitochondria; therefore, we ran CAT a second time using `--chain-mode` with the hg38ToPanTro6.over.chain.gz file obtained from the UCSC genome browser to obtain mitochondrial genome annotations for chimpanzee. We then combined the autosomal and mitochondrial alignments to produce a single chimpanzee annotation set. We then filtered CAT-based annotations to exclude features that were not shared by both species in each pairwise comparison. We further filtered the data to account for differences in transcript length (see **Table** S3).

### Orthologous exons

We previously used BLAT (51) to generate lists of orthologous exons with high sequence identity that reciprocally map to the reference genome of each species (7,32,34,35,52). Here, instead of using BLAT, we built orthologous exon annotations by filtering exons from the CAT genome annotations, removing exons with less than 85% sequence identity or that differed by more than 10% in sequence length (**Table** S3). This approach allowed for a straightforward comparison between ‘annotation transfer’, ‘orthologous exons’, and CrossFilt.

### Differential expression analysis

We performed DE analysis separately in each tissue and species comparison using a standard pipeline. We restricted our analysis to genes that had at least one count in at least one sample from both species (either human/chimpanzee or human/macaque). We further filtered out gene sets using default parameters from the `filterByExpr` function in edgeR (41,42). This function keeps genes that meet a counts-per-million (**CPM**) threshold in a number of samples equal to the smallest group size. We calculated library sizes and standardized the data using trimmed-mean of M-values (53) and generated log-transformed CPM and precision weights using the `voom` function in edgeR (54). We used these CPM values for differential expression analysis, except in the ‘dual reference’ approach. Here, we generated log-transformed transcripts-per-million (**TPM**) in place of CPM to account for gene length differences. Using the limma function `lmFit` (55), we fit a linear model using a single fixed effect for species and moderated the t-statistics using the `eBayes` functions in limma. To rule out normalization as a potential factor explaining differences between the ‘dual reference’ approach and the other pipelines, we repeated these steps using TPM normalization in all of the approaches (**Figure S4**).

### Sharing of differentially expressed genes between tissues

We used *mash* (38) to estimate sharing of DE effects between tissues in each species separately. Mash takes the DE effect sizes and standard errors as input. We used effect size estimates from limma and computed standard errors from the effect sizes and p-values. We ran *mash* using both canonical and data-driven covariance matrices, using four principal components for the latter. We first checked for sharing of effects between each pair of tissues. We considered a gene to have shared differential expression if it had a false sign rate below 5% in at least one tissue, and if posterior estimates of the effect sizes were within a factor of 1.5 of each other in all pairwise tissue comparisons.

### Gene set enrichment analysis

We performed gene set enrichment analysis using the R package fgsea (56). We obtained Gene Ontology biological process gene sets from the Molecular Signature Database v7.5.1. We used the ranked t-statistics from each tissue-by-species comparison to perform GSEA for each biological process.

## Declarations

### Ethics approval and consent to participate

Not applicable.

### Consent for publication

Not applicable.

## Availability of data and materials

All data used in this work are publicly available. The methods we have implemented are available in github repositories https://github.com/kennethabarr/CrossFilt and https://github.com/kennethabarr/ConsensusGenomeTools.

## Funding

This work was supported by NIH grants R35GM131726 and R01HG010772, to YG.

### Author contributions

KB conceptualized the study, designed the method, generated results, and performed the analysis. YG provided funding for the project and wrote the manuscript, with input and edits from KB.

## Acknowledgements

We thank reviewer 2 for their feedback on our previous paper (35). Their questions regarding the influence of alignment bias inspired this work. We thank Genevieve Housman for helpful discussion. We thank the creators of CrossMap (57), whose implementation of liftOver helped us get started with this project. We also acknowledge Natalia Gonzales for her assistance writing and revising the manuscript.

## Supplementary Figures and Tables

**Table S1:** Annotation summary. Number of genes, transcripts, and genome coverage, expressed bases covered by exons, as well as percent of the genome covered by exons.

| Annotation | Human / Chimpanzee |  |  |  | Human / Rhesus |  |  |  |
| --- | --- | --- | --- | --- | --- | --- | --- | --- |
|  | Consensus /<br>Single-reference | Dual-reference | Orthologous exons | Annotation Transfer | Consensus /<br>Single-reference | Dual-reference | Orthologous exons | Annotation Transfer |
| Genes | 38,584 | 17,443 | 36,102 | 37,960 | 38,584 | 16,885 | 34,694 | 37,819 |
| Protein coding genes | 19,405 | 17,416 | 18,473 | 19,177 | 19,405 | 16,855 | 18,238 | 19,123 |
| Human transcripts | 226,006 | 143,567 | 215,824 | 215,512 | 226,006 | 157,225 | 211,969 | 201,363 |
| Non-human transcripts | 226,006 | 77,605 | 215,766 | 215,512 | 226,006 | 63,187 | 211,847 | 201,363 |
| Human Coverage (Mbp) | 146.4 | 96.1 | 134.8 | 140.6 | 146.4 | 101.2 | 116.4 | 129.7 |
| Non-human Coverage (Mbp) | 146.4 | 76.2 | 134.7 | 140.0 | 146.4 | 67.1 | 115.9 | 128.2 |
| Human Coverage (%) | 4.72 | 3.10 | 4.35 | 4.53 | 4.72 | 3.26 | 3.76 | 4.19 |
| Non-human Coverage (%) | 4.72 | 2.50 | 4.42 | 4.59 | 4.72 | 2.26 | 3.90 | 4.31 |

**Table S2:**
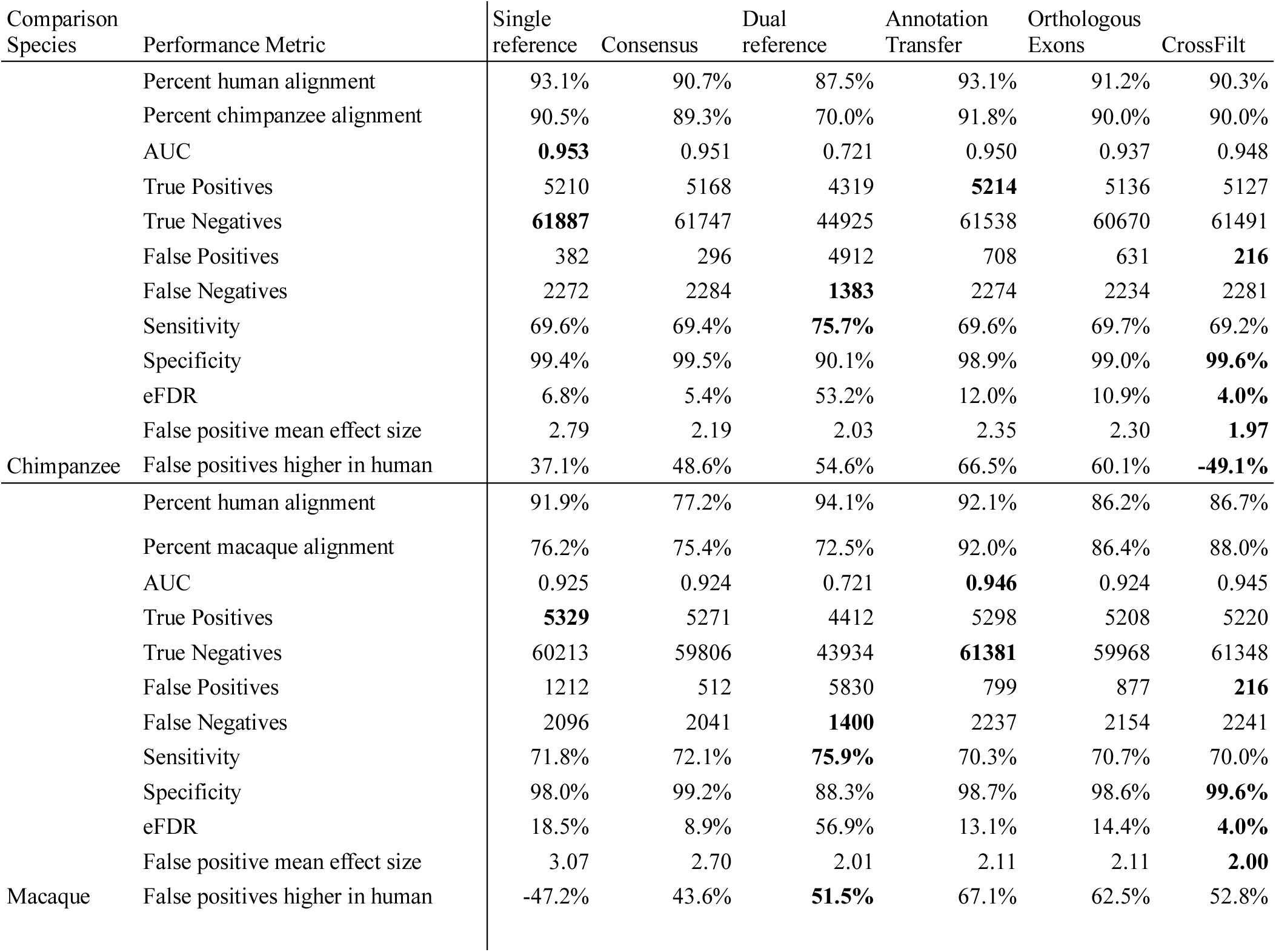
Performance metrics on simulated reads. To allow performance comparisons across species we computed performance metrics using the subset of genes that were tested in both species comparisons. The ‘percent alignment’ is the percent of simulated reads that were aligned and counted for the gene subset in each species. ‘False positives higher in human’ shows the percentage of false positives that have higher expression in humans relative to the comparison species.

| Comparison Species | Performance Metric | Single reference | Consensus | Dual reference | Annotation Transfer | Orthologous Exons | CrossFilt |
| --- | --- | --- | --- | --- | --- | --- | --- |
| Chimpanzee | Percent human alignment | 93.1% | 90.7% | 87.5% | 93.1% | 91.2% | 90.3% |
|  | Percent chimpanzee alignment | 90.5% | 89.3% | 70.0% | 91.8% | 90.0% | 90.0% |
|  | AUC | **0.953** | 0.951 | 0.721 | 0.950 | 0.937 | 0.948 |
|  | True Positives | 5210 | 5168 | 4319 | **5214** | 5136 | 5127 |
|  | True Negatives | **61887** | 61747 | 44925 | 61538 | 60670 | 61491 |
|  | False Positives | 382 | 296 | 4912 | 708 | 631 | **216** |
|  | False Negatives | 2272 | 2284 | **1383** | 2274 | 2234 | 2281 |
|  | Sensitivity | 69.6% | 69.4% | **75.7%** | 69.6% | 69.7% | 69.2% |
|  | Specificity | 99.4% | 99.5% | 90.1% | 98.9% | 99.0% | **99.6%** |
|  | eFDR | 6.8% | 5.4% | 53.2% | 12.0% | 10.9% | **4.0%** |
|  | False positive mean effect size | 2.79 | 2.19 | 2.03 | 2.35 | 2.30 | **1.97** |
|  | False positives higher in human | 37.1% | 48.6% | 54.6% | 66.5% | 60.1% | **-49.1%** |
| Macaque | Percent human alignment | 91.9% | 77.2% | 94.1% | 92.1% | 86.2% | 86.7% |
|  | Percent macaque alignment | 76.2% | 75.4% | 72.5% | 92.0% | 86.4% | 88.0% |
|  | AUC | 0.925 | 0.924 | 0.721 | **0.946** | 0.924 | 0.945 |
|  | True Positives | **5329** | 5271 | 4412 | 5298 | 5208 | 5220 |
|  | True Negatives | 60213 | 59806 | 43934 | **61381** | 59968 | 61348 |
|  | False Positives | 1212 | 512 | 5830 | 799 | 877 | **216** |
|  | False Negatives | 2096 | 2041 | **1400** | 2237 | 2154 | 2241 |
|  | Sensitivity | 71.8% | 72.1% | **75.9%** | 70.3% | 70.7% | 70.0% |
|  | Specificity | 98.0% | 99.2% | 88.3% | 98.7% | 98.6% | **99.6%** |
|  | eFDR | 18.5% | 8.9% | 56.9% | 13.1% | 14.4% | **4.0%** |
|  | False positive mean effect size | 3.07 | 2.70 | 2.01 | 2.11 | 2.11 | **2.00** |
|  | False positives higher in human | -47.2% | 43.6% | **51.5%** | 67.1% | 62.5% | 52.8% |

**Figure S1:**
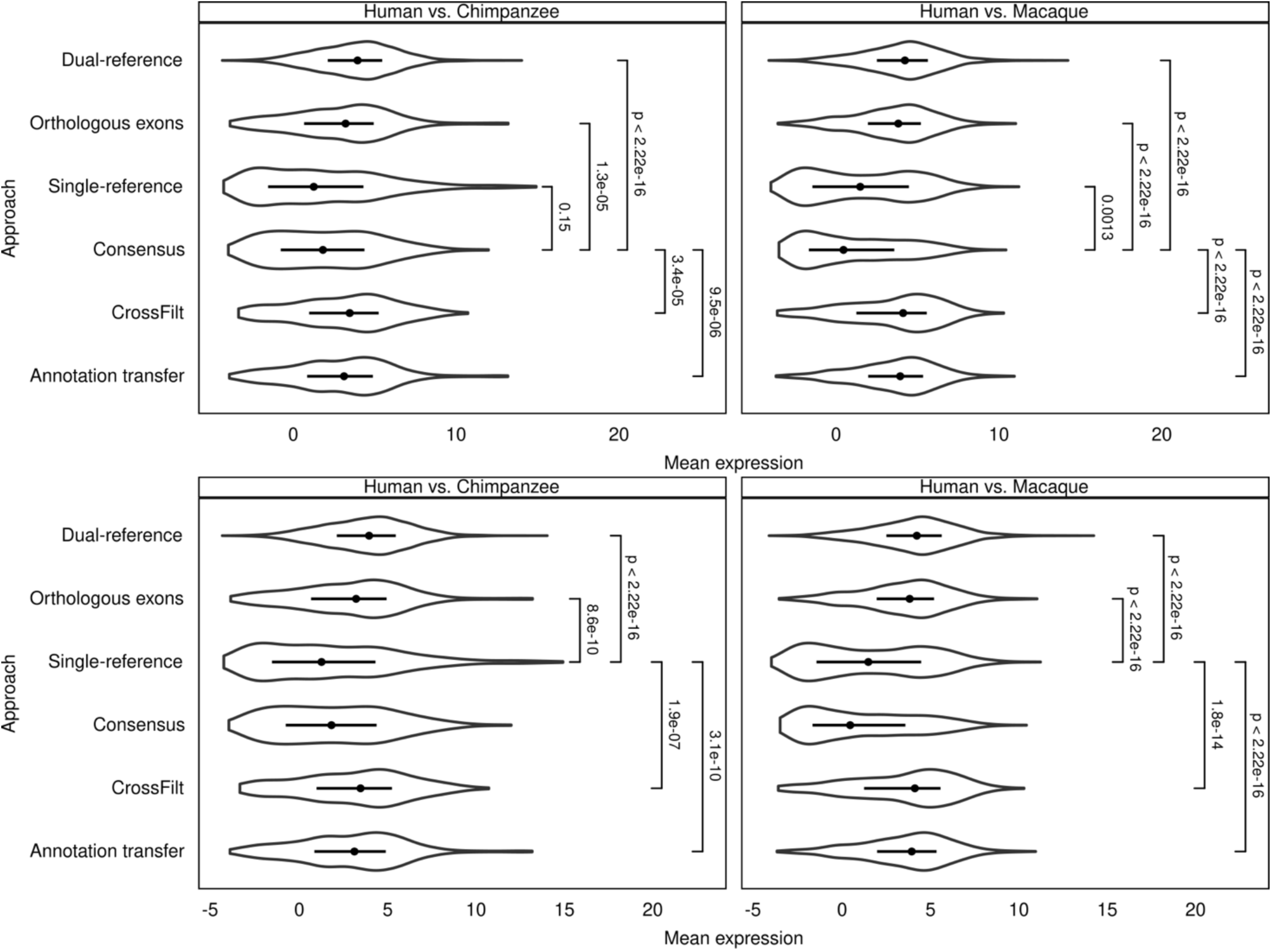
Mean expression of false positives. We compared mean expression levels for false positive at a 5% FDR in the ‘consensus’ (Top) and ‘single-reference’ (Bottom) to false positives from all other approaches. Wilcoxon rank sum test p-values are indicated.

**Figure S2:**
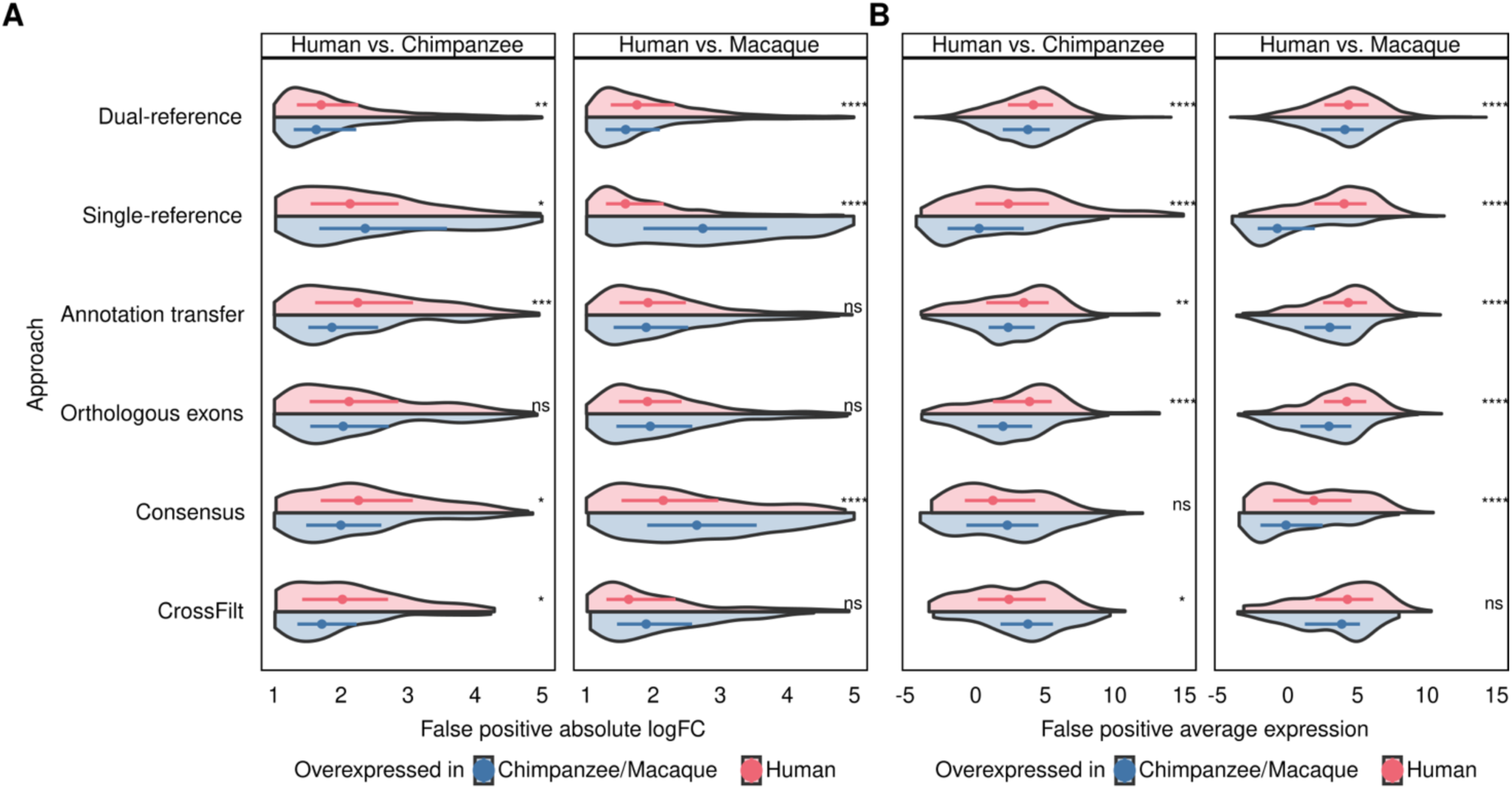
Properties of false positives by species. **(A)** Absolute difference in expression levels for false positives, split by the species in which the gene has higher expression. **(B)** Average expression levels for false positives, split by the species in which the gene has higher expression. P-values were computed using Wilcoxon rank-sum (ns = not significant; * p ≤ 0.05; ** p ≤ 0.01; *** p ≤ 0.001; **** p ≤ 0.0001).

**Figure S3:**
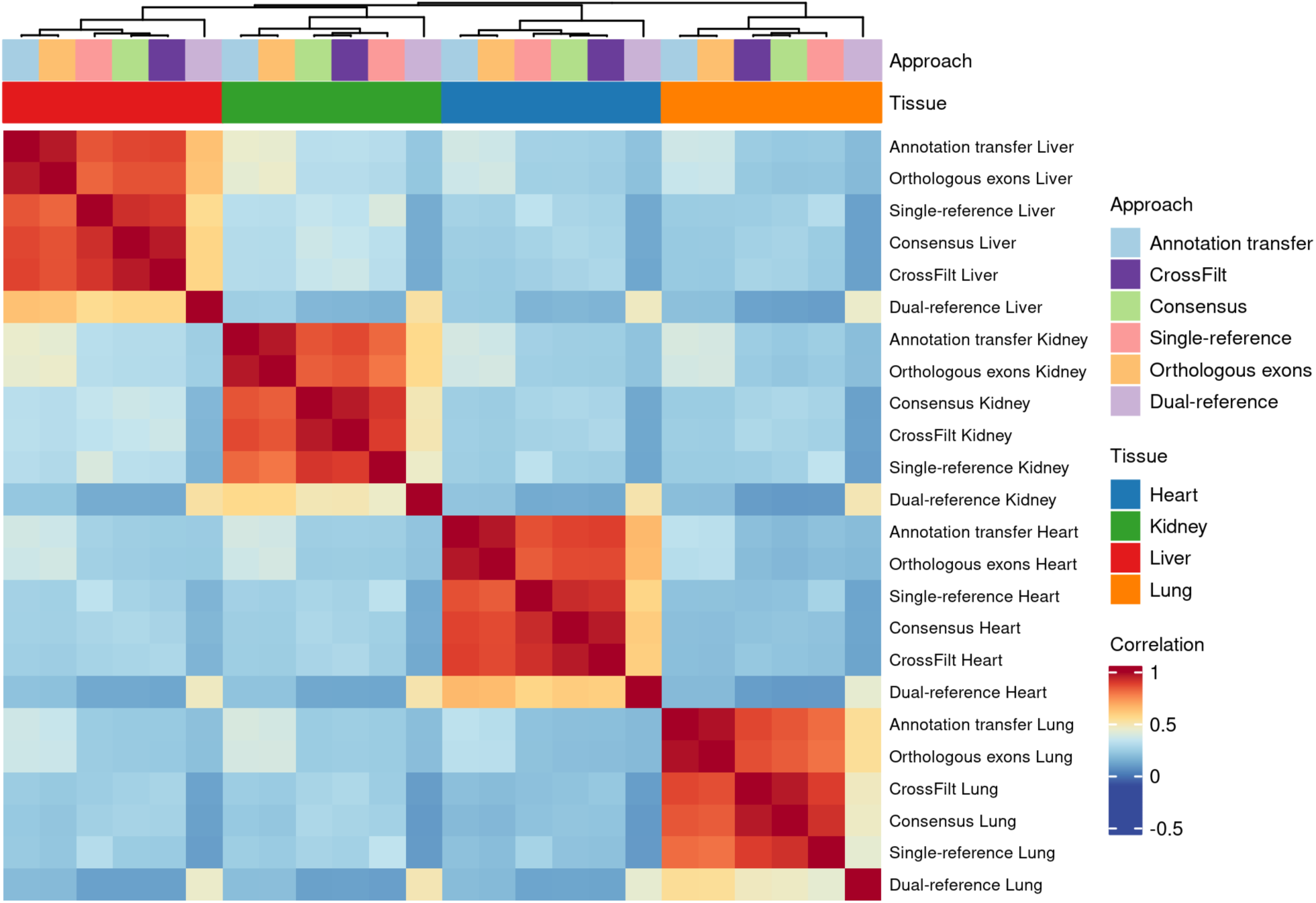
Heatmap of the Pearson correlation in human-macaque effect size estimates for genes detected as DE in at least approach and tissue. Order of rows and columns is set by hierarchical clustering.

**Figure S4:**
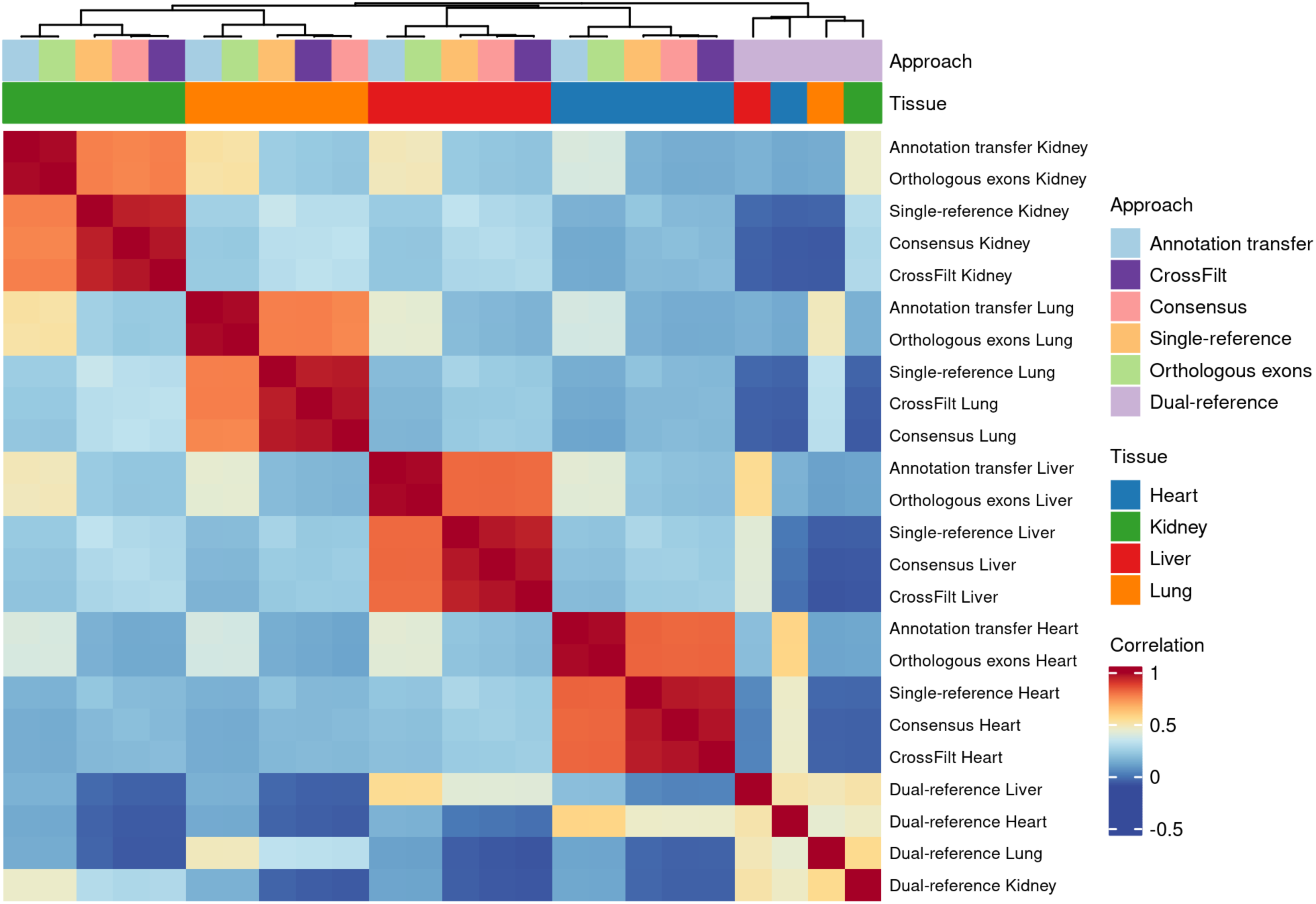
Regeneration of Figure 3A using TPM normalization on all approaches instead of CPM normalization.

**Table S3:** Genomes and annotations used.

| Alignment | Species | Comparison | Annotation | Genome |
| --- | --- | --- | --- | --- |
| Dual-reference | Human | Human vs. Chimpanzee | 2024 10x Cellranger, filtered for common names | hg38 |
| Dual-reference | Chimpanzee | Human vs. Chimpanzee | panTro6 refSeq, filtered for common names | panTro6 |
| Dual-reference | Human | Human vs. Rhesus | 2024 10x Cellranger, filtered for common names | hg38 |
| Dual-reference | Rhesus | Human vs. Rhesus | rheMac10 refSeq, filtered for common names | rheMac10 |
| Single-reference | Human | All | 2024 10x Cellranger | hg38 |
| Single-reference | Chimpanzee | All | 2024 10x Cellranger | hg38 |
| Single-reference | Rhesus | All | 2024 10x Cellranger | hg38 |
| Consensus | Human | Human vs. Chimpanzee | 2024 10x Cellranger | hg38/panTro6 consensus genome |
| Consensus | Chimpanzee | Human vs. Chimpanzee | 2024 10x Cellranger | hg38/panTro6 consensus genome |
| Consensus | Human | Human vs. Macaque | 2024 10x Cellranger | hg38/rheMac10 consensus genome |
| Consensus | Macaque | Human vs. Macaque | 2024 10x Cellranger | hg38/rheMac10 consensus genome |
| Annotation transfer | Human | Human vs. Chimpanzee | 2024 10x Cellranger annotation, filtered for common transcripts with less than 10% length difference with chimpanzee | hg38 |
| Annotation transfer | Chimpanzee | Human vs. Chimpanzee | CAT transferred annotation, filtered for common transcripts with less than 10% length difference with human | panTro6 |
| Annotation transfer | Human | Human vs. Macaque | 2024 10x Cellranger annotation, filtered for common transcripts with less than 10% length difference with macaque | hg38 |
| Annotation transfer | Macaque | Human vs. Macaque | CAT transferred annotation, filtered for common transcripts with less than 10% length difference with human | rheMac10 |
| Orthologous exon | Human | Human vs. Chimpanzee | 2024 10x Cellranger annotation, filtered for common exons with greater than 85% sequence identity and less than 10% length difference in chimpanzee | hg38 |
| Orthologous exon | Chimpanzee | Human vs. Chimpanzee | CAT transferred annotation, filtered for common exons with greater than 85% sequence identity and less than 10% length difference in human | panTro6 |
| Orthologous exon | Human | Human vs. Macaque | 2024 10x Cellranger annotation, filtered for common exons with greater than 85% sequence identity and less than 10% length difference in rhesus | hg38 |
| Orthologous exon | Macaque | Human vs. Macaque | CAT transferred annotation, filtered for common exons with greater than 85% sequence identity and less than 10% length difference in rhesus | rheMac10 |
| CrossFilt | Human | All | 2024 10x Cellranger | hg38 |
| CrossFilt | Chimpanzee | All | Cat transferred annotation | panTro6 |
| CrossFilt | Macaque | All | Cat transferred annotation | rheMac10 |

